# AI-Driven Design of Nanobinders Targeting the TSLPR Heterodimer Interface to Suppress Type 2 Inflammatory Signaling

**DOI:** 10.64898/2026.07.20.739581

**Authors:** Woo-Chan Ahn, Min-Jung Son, Jae-Rin Kim, Seong-Ryeong Go, Ju-Eun Kang, Junsoo Park, Young-Hoon Lee, Su-Jin Lee, Jeong-Hoon Kim, Juyeon Jung, Eui-Jeon Woo, Young-Joo Jeon, Kwang-Hyun Park

**Affiliations:** Bio-design & Editing Research Center, Korea Research Institute of Bioscience & Biotechnology (KRIBB), Daejeon 34141, Republic of Korea; Metabolic Regulation Research Center, KRIBB, Daejeon 34141, Republic of Korea; Department of Functional Genomics, KRIBB School of Bioscience, University of Science and Technology (UST), Daejeon 34113, Republic of Korea; Critical Diseases Diagnostics Convergence Research Center, KRIBB, Daejeon 34141, Republic of Korea; Department of Proteome Structural Biology, KRIBB School of Bioscience, UST, Daejeon 34113, Republic of Korea; Orphan Disease Therapeutic Target Research Center, KRIBB, Daejeon 34141, Republic of Korea; Bionanotechnology Research Center, KRIBB, Daejeon 34141, Republic of Korea

**Keywords:** protein design, antagonist, nanobinder, interface-mimetic grafting, TSLP

## Abstract

Aberrant thymic stromal lymphopoietin (TSLP) signaling is a central driver of type 2 inflammatory diseases, yet the only approved TSLP-targeted therapy is a 150 kDa monoclonal antibody whose bulky format limits tissue penetration and precludes inhaled delivery. Here, we report an AI-driven framework for designing ultra-compact *de novo* nanobinders that suppress TSLP signaling by sterically disrupting assembly of the TSLPR-IL-7Rα heterodimer. We compare two structure-guided strategies, namely purely *de novo* helical bundle generation and interface-mimetic grafting of native binding motifs onto designed scaffolds. Although both yield nanomolar binders, only orthosteric mimicry of the native cytokine geometry blocks receptor heterodimerization, showing that functional antagonism is governed by precise epitope geometry rather than affinity alone. After library-based maturation, the lead nanobinder TRB5.1 is a hyper-stable monomer (T_m_ ∼97.4 °C) with single-digit nanomolar affinity (K_D_ = 9.7 nM) and strict selectivity over related α-chain interleukin receptors. TRB5.1 suppresses TSLP-induced JAK1 and STAT5 phosphorylation across multiple cellular models and drives a transcriptome-wide reversal of the pathogenic type 2 program. This work delivers a developable, potentially inhalable non-antibody lead and a scalable blueprint for antagonizing heterodimeric cytokine receptors.

## Introduction

Asthma, atopic dermatitis, and other type 2 inflammatory diseases together affect hundreds of millions of patients worldwide and impose a major socioeconomic burden [1]. These conditions share a common driver, namely abnormal activation of the epithelial alarmin and T-helper 2 (Th2) cytokine network [2]. Thymic stromal lymphopoietin (TSLP), released from airway and skin epithelial cells in response to allergens, viruses, pollutants, and oxidative stress, activates dendritic cells, mast cells, and group 2 innate lymphoid cells, triggering downstream production of interleukin (IL)-4, IL-5, IL-9 and IL-13 [2–4]. Because TSLP acts as the master switch of this network, it has emerged as one of the most attractive therapeutic targets in immunology [5, 6].

The only clinically validated therapy targeting this axis is tezepelumab, an anti-TSLP monoclonal antibody that neutralizes soluble TSLP before it engages its receptor and is approved for severe uncontrolled asthma [7, 8]. Its full-length immunoglobulin G (IgG) format, however, illustrates the typical limitations of conventional antibody biologics [9, 10]. This 150 kDa size restricts mucosal and tissue penetration, confining administration mostly to intravenous or subcutaneous injection and ruling out patient-friendly routes such as inhalation or topical application [11]. Mammalian cell-based production, strict cold-chain logistics, and Fc-mediated immune effects further drive up costs and limit broader clinical use [12, 13]. TSLP signals through a heterodimer of the TSLP receptor (TSLPR) and the IL-7 receptor α chain (IL-7Rα), activating STAT5 in diverse hematopoietic cells. This heterodimer provides an alternative point of intervention on the receptor side (**Fig. 1a)**.

**Figure 1.**
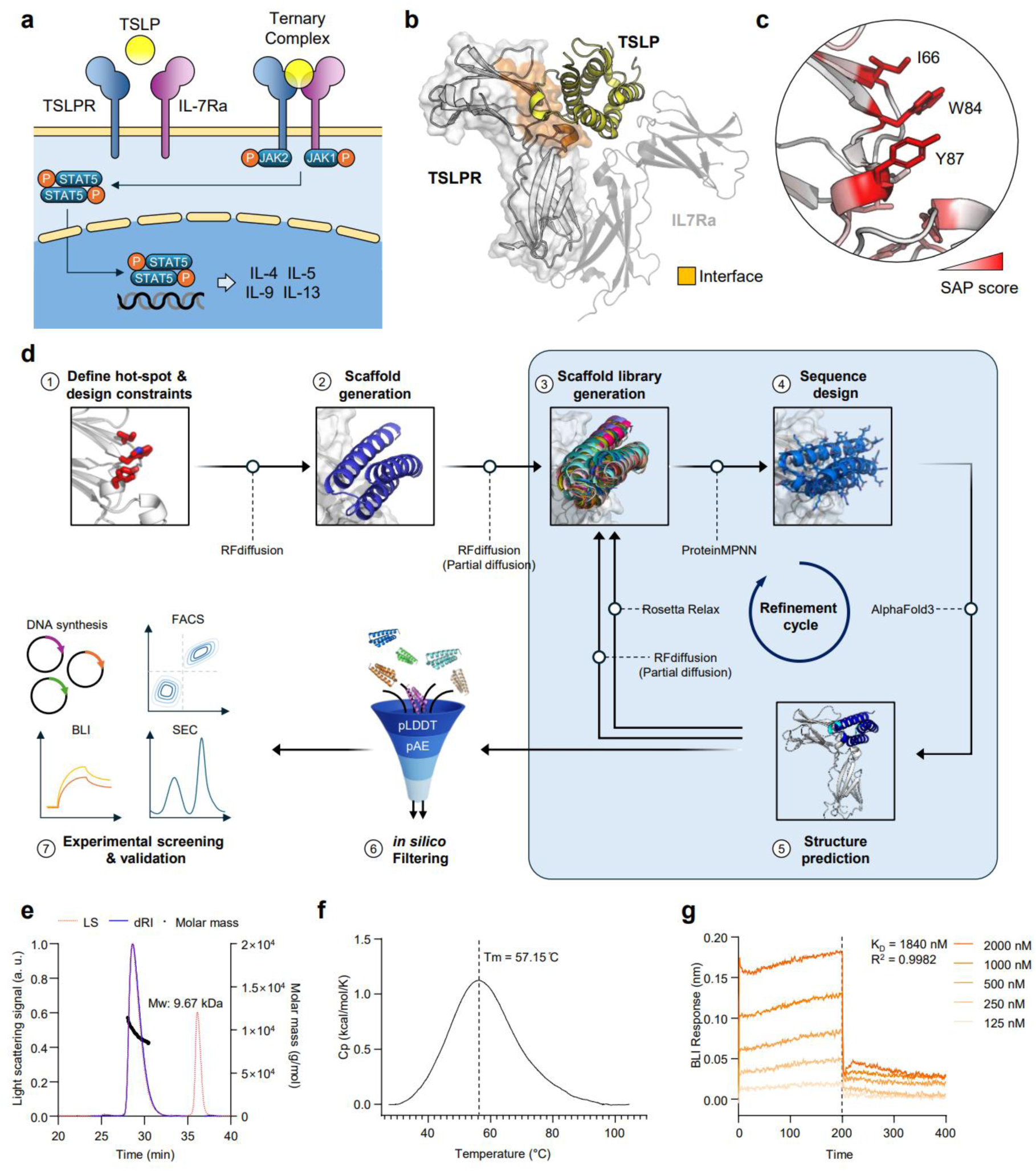
Design workflow and initial characterization of *de novo* helical nanobinders targeting TSLPR. (a) TSLP-mediated receptor assembly and signaling. TSLP bridges TSLPR and IL-7Rα at the cell surface, driving intracellular JAK-STAT signaling and downstream cytokine expression. (b) Structure of the human TSLP-TSLPR-IL-7Rα ternary complex. The interface between TSLP and TSLPR used for binder design is shown, with the receptor-binding surface on TSLPR highlighted in yellow. (c) Close-up of the TSLPR hot-spot region. Residues are colored by SAP score, and the selected hydrophobic residues I66, W84, and Y87 are shown as sticks. (d) Computational and experimental design workflow. The pipeline comprises hot-spot and design-constraint definition, scaffold generation, scaffold library generation, sequence design, structure prediction, *in silico* filtering, and experimental screening and validation. (e) SEC-MALS profile of purified TRB2.0. Light scattering, differential refractive index, and calculated molar mass traces are shown. (f) DSC thermogram of purified TRB2.0. A single cooperative unfolding transition is observed. (g) BLI sensorgrams of TRB2.0 binding to TSLPR. Dose-dependent responses across a concentration series are shown.

Artificial intelligence (AI) has reshaped protein engineering [14, 15]. RFdiffusion designs new backbones for a target surface [16], ProteinMPNN assigns sequences that fold stably onto them [17], and AlphaFold3 (AF3) predicts the complexes with near-experimental accuracy [18]. This allows reliable computational filtering before any wet-lab work. Together these tools yield ultra-small *de novo* miniproteins of roughly 5 to 10 kDa that bind targets with atomic-level precision, a class we term nanobinders [16, 19, 20]. Although nanobinders resemble camelid nanobodies, they differ fundamentally in origin and developability. Nanobodies arise from immunization or library screening and are confined to the immunoglobulin fold [21]. In contrast, nanobinders are designed entirely *in silico* with custom shapes, offering better tissue penetration, low-cost microbial production, programmable stability, and reduced immune risk [22, 23].

Blocking a heterodimerization interface requires binders that occupy the exact contact residues used by the natural cytokine [24, 25]. This is a challenge that unconstrained generative sampling alone cannot reliably solve. Building on a careful structural analysis of the TSLP-TSLPR-IL-7Rα ternary complex, we employed two complementary structure-guided strategies within a unified pipeline [6]. The first strategy, *de novo* helical bundle targeting, used RFdiffusion to scaffold a helical bundle that displays designed hot-spot residues against the cytokine-binding cleft of TSLPR. This produced entirely new topologies that are not constrained by the natural cytokine fold. The second strategy, interface-mimetic stabilization and maturation, took the TSLP-derived binding region and grafted it onto a stabilizing scaffold. This grafted construct was then refined by repeated computational maturation to improve folding stability, surface complementarity, and binding energy, while keeping the natural binding geometry intact.

Computational candidates from both strategies were displayed on the yeast surface and screened by fluorescence-activated cell sorting (FACS) using fluorescently labeled TSLPR. Top binders from each strategy were then advanced through library-based affinity maturation under increasingly strict sorting. The resulting nanobinders were evaluated across three levels. We measured expression yield, oligomeric state, and thermal stability, quantified binding affinity and specificity against related interleukin receptors, and assessed blockade of TSLP-induced STAT5 signaling. When the two strategies were compared side by side, the interface-mimetic strategy produced nanobinders with higher affinity and greater biophysical robustness than the *de novo* helical bundle approach. This indicates that the native binding geometry offers a clear advantage for antagonizing challenging targets such as cytokine receptors. Ultimately, this structure-guided computational framework provides a scalable blueprint for developing next-generation, non-antibody biologics.

## Results

### *De Novo* Design of Helical Nanobinders Targeting TSLPR

To develop nanobinders capable of antagonizing TSLPR signaling, we first analyzed the structural basis of natural TSLP recognition using the TSLP-TSLPR-IL-7Rα ternary complex (PDB 5J11) as a reference [6]. In this assembly, TSLP is sandwiched between the two receptors, engaging TSLPR across a broad, discontinuous interface that spans multiple secondary structural elements **(Fig. 1b)**. To translate this topologically complex interface into a tractable design objective, we sought to isolate the contact residues that dominate the natural interaction. Spatial Aggregation Propensity (SAP) scoring of the receptor interface revealed a hydrophobic hot-spot centered on the conserved TSLPR residues I66, W84, and Y87 **(Fig. 1c)**. This localized patch defined the spatial pharmacophore that any *de novo* binder must engage to compete with endogenous TSLP.

To engineer compact nanobinders that occlude this cytokine-binding cleft without inheriting the topological constraints of the native cytokine fold, we implemented an iterative, generative AI-driven pipeline **(Fig. 1d)**. Initial helical scaffolds targeting the hot-spot were generated by AI-based backbone design, with the I66/W84/Y87 patch supplied as a spatial constraint. This constraint steered the generated backbones to dock against the target cleft rather than at off-target sites on the receptor surface. Sequence design was then applied to assign amino acid identities optimized for both folding stability and shape complementarity with the receptor interface. To refine the binding geometry at near-atomic resolution, we performed an additional round of localized backbone remodeling, followed by sequence redesign and structure modeling for validation. The designed models aligned closely with their AF3-predicted topologies, with the engineered interface residues exhibiting high structural confidence **(Fig. S1)**. To prioritize candidates with the highest probability of folding and target engagement, we applied in silico filtering thresholds. A refined pool of 24 designs meeting the criteria of monomeric folding confidence (pLDDT) and interchain predicted aligned error (pAE) was retained for experimental validation **(Fig. S2a)**.

Translating these *in silico* predictions to *in vitro* validation, the computationally prioritized candidates were displayed on the yeast surface and subjected to screening by FACS against fluorescently labeled TSLPR. Successive rounds of sorting produced a progressive enrichment of the TSLPR-binding population, with the target-gated fraction expanding from less than 10% in the initial round to approximately 45% after iterative selection **(Fig. S2b)**. This enrichment trajectory indicated that a substantial proportion of the designed library possessed target-specific binding activity rather than non-specific surface interactions.

To evaluate the biophysical properties of the *de novo* designs, a representative binder, TRB2.0, was subcloned into a microbial expression vector, expressed in *E. coli*, and purified as a highly soluble protein in quantities sufficient for biophysical characterization. Size-exclusion chromatography coupled with multi-angle light scattering (SEC-MALS) confirmed its biophysical homogeneity. The protein eluted as a single monodisperse peak at its theoretical monomeric mass, with no detectable higher-order oligomers or aggregates **(Fig. 1e)**. Thermal stability analysis by differential scanning calorimetry (DSC) revealed a cooperative unfolding transition with a melting temperature (T_m_) of 57.15 °C, confirming that the designed helical bundle adopts a well-defined and thermodynamically stable structure **(Fig. 1f)**. Bio-layer interferometry (BLI) showed dose-dependent binding to immobilized TSLPR, with a calculated equilibrium dissociation constant (K_D_) of approximately 1,840 nM **(Fig. 1g)**. Collectively, these results confirm that a purely structure-guided *de novo* strategy can deliver a stable, monomeric nanobinder targeting the TSLPR cleft. Importantly, this approach establishes a robust structural foundation that primes the candidate for subsequent affinity maturation.

### Interface-Mimetic Design of Motif-Grafted Nanobinders Targeting TSLPR

In parallel with the *de novo* helical bundle approach, we employed a second structure-guided strategy that directly leverages the natural binding geometry. Structural analysis of native TSLP revealed an atypical recognition mode in which the TSLPR-binding motif consists not of a single continuous region but of a binding α-helix (α4) and the binding loop that together contact the receptor surface **(Fig. 2a)**. Although this loop contributes directly to binding, it also exposes hydrophobic residues, increases conformational entropy, and is prone to proteolytic degradation, compromising thermodynamic stability and developability. To preserve the natural contact residues while eliminating this weakness, we computationally extracted the α-helix (residues 139–159) and the binding loop (residues 53–74). These elements were grafted onto a de novo designed three-helix bundle scaffold, with the non-binding disordered loop removed entirely **(Fig. 2a and 2b)**. The resulting topology retains the natural binding face of TSLP while replacing the disordered loop with a rigid designed backbone, integrating natural specificity and designed stability into a single compact construct **(Fig. 2c)**.

**Figure 2.**
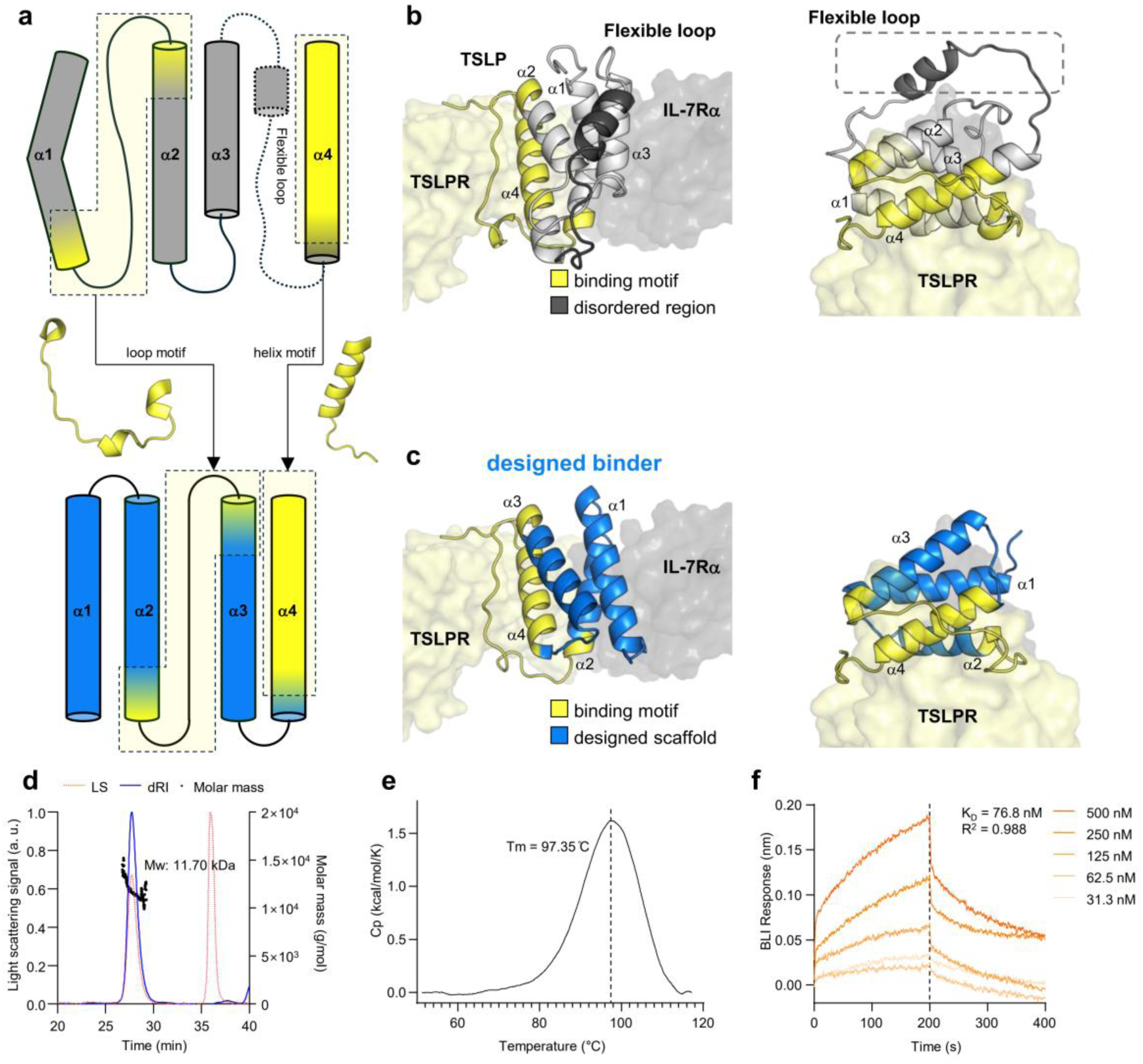
Interface-mimetic design of scaffolded TSLPR-binding nanobinders. (a) Interface-mimetic design strategy. Native TSLP-derived TSLPR-binding elements are shown together with the flexible region selected for replacement by a designed scaffold. (b) Native TSLP-TSLPR interaction. The TSLP-derived binding motif and disordered region are indicated on the receptor-bound structure. (c) Designed binder bound to TSLPR. The retained binding motif and designed scaffold are shown on the TSLPR-binding surface. (d) SEC-MALS profile of purified TRB5.0. Light scattering, differential refractive index, and calculated molar mass traces are shown. (e) DSC thermogram of purified TRB5.0. A single cooperative unfolding transition is observed. (f) BLI sensorgrams of TRB5.0 binding to TSLPR. Dose-dependent responses across a concentration series are shown.

Starting from this design, we applied iterative cycles of backbone generation, sequence design, and structure modeling to progressively optimize the grafted motif and surrounding scaffold. This refinement yielded three representative backbone topologies that stably presented the natural binding face. Using pLDDT and pAE as primary filters, we then generated 28 sequence variants distributed across these three backbones **(Fig. S3a)**. All 28 designs were displayed on the yeast surface and screened against fluorescently labeled TSLPR by FACS. Successive rounds progressively enriched the target-binding population, and sequence analysis of the enriched clones identified three distinct variants sharing the same backbone topology but carrying different surface sequences **(Fig. S3b and S4a)**. Structural alignment of the three variants with native TSLP confirmed that the engineered nanobinders closely reproduce the natural binding geometry while adopting a more compact and rigid global topology **(Fig. S4b)**. The conserved predicted interface reconstructs the polar and hydrophobic interaction networks with the TSLPR hot-spot residues I66, W84, and Y87, using the same contact residues as native TSLP **(Fig. S4c)**. The recognition logic of the natural TSLP-TSLPR interaction is thus preserved on a designed scaffold free of the disordered loop.

For experimental validation, the three selected variants were produced in *E. coli*. All three were expressed at high yield, but two of them underwent marked degradation during purification. The remaining variant, which was purified to homogeneity without detectable degradation, was selected as the lead and designated TRB5.0. SEC-MALS confirmed that TRB5.0 eluted as a single monodisperse peak at its theoretical monomeric mass, with no detectable higher-order species **(Fig. 2d)**. Replacing the disordered loop with a designed continuous backbone translated into a marked gain in thermal stability. DSC revealed a cooperative unfolding transition with a T_m_ of 97.35 °C **(Fig. 2e)**. This is approximately 40 °C higher than the ∼57 °C T_m_ of the *de novo* helical binder, well within the range favorable for therapeutic developability. BLI showed dose-dependent binding with slow dissociation under buffer flow, yielding an equilibrium dissociation constant (K_D_) of approximately 76.8 nM, roughly a 24-fold improvement over the ∼1,840 nM K_D_ of the *de novo* binder **(Fig. 2f)**. Collectively, these results show that anchoring the natural binding geometry on a rigid designed scaffold integrates binding fidelity and thermodynamic robustness into a single compact construct that surpasses the *de novo* helical bundle baseline in both affinity and thermal stability. With both baseline binders validated, the top candidates from each strategy were advanced to library-based affinity maturation.

### Library-Based Affinity Maturation and Specificity Assessment

To enhance the binding affinity of the baseline nanobinders and strengthen their antagonistic potency against TSLPR, we performed library-based affinity maturation. For each baseline nanobinder, we constructed a site-saturation mutagenesis (SSM) library in which every interface position was independently substituted with all 20 amino acids. Each library was then matured by yeast surface display, with iterative rounds of FACS performed under progressively stringent conditions achieved by stepwise reduction of the fluorescently labeled TSLPR concentration. In both libraries, a pronounced shift from the baseline distribution toward variant populations with enhanced binding activity was observed **(Fig. 3a and 3b)**.

**Figure 3.**
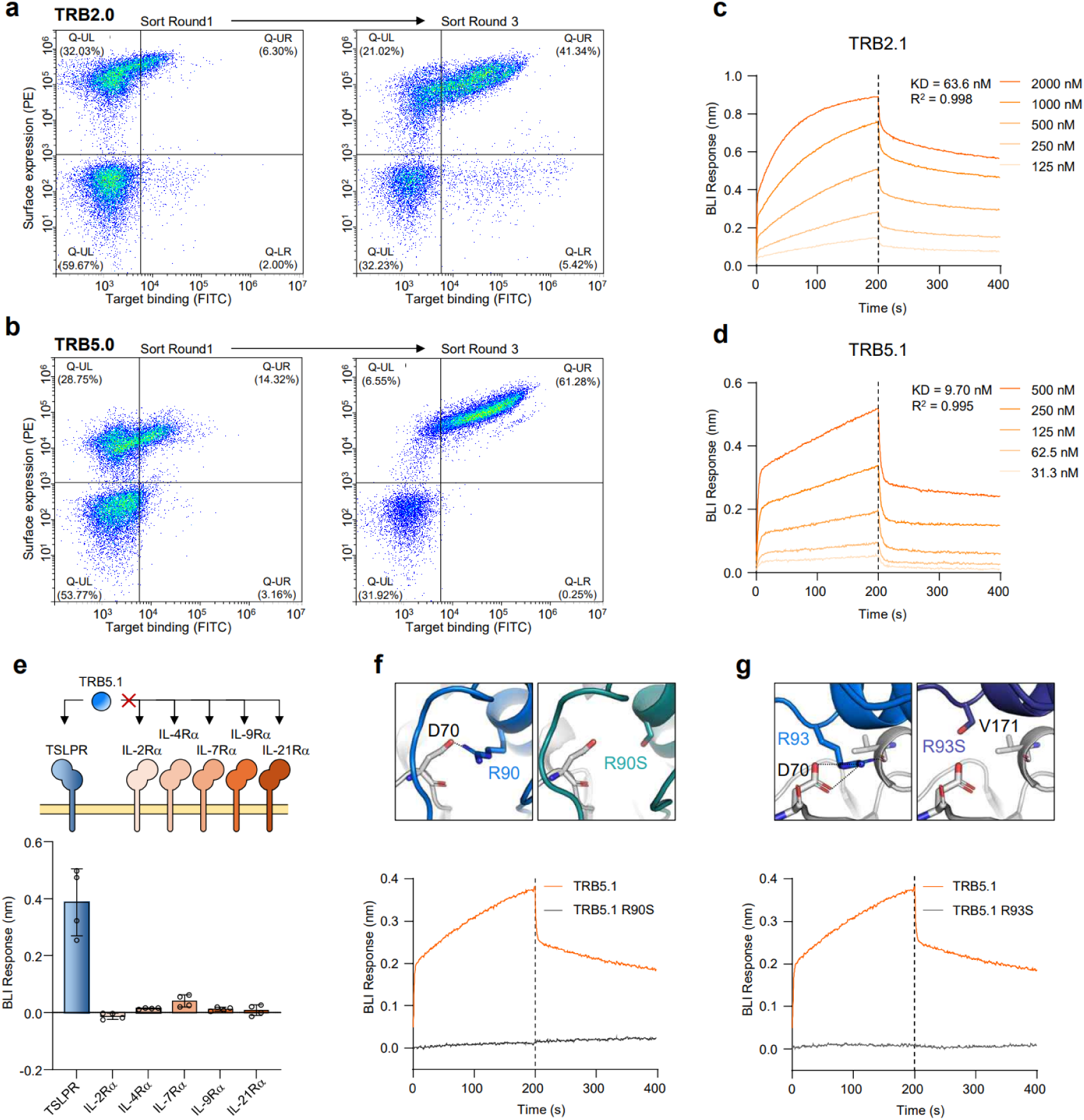
Affinity maturation and specificity assessment of TSLPR nanobinders. (a) FACS plots of the TRB2.0 yeast library before and after affinity maturation. The x-axis shows target binding (FITC) and the y-axis shows surface expression (PE). Gate percentages indicate the proportion of double-positive cells in each round. (b) FACS plots of the TRB5.0 yeast library before and after affinity maturation. The x-axis shows target binding (FITC) and the y-axis shows surface expression (PE). Gate percentages indicate the proportion of double-positive cells in each round. (c) BLI sensorgrams of TRB2.1 binding to TSLPR. Responses are shown at the indicated concentrations. (d) BLI sensorgrams of TRB5.1 binding to TSLPR. Responses are shown at the indicated concentrations. (e) Binding specificity of TRB5.1 across the cytokine receptor α-chain family. The schematic indicates the receptor panel tested, and the bar graph summarizes binding responses against TSLPR and five related receptors. Each dot represents one of four independent replicates, and bars indicate mean ± SD. (f) Role of R90 at the predicted TRB5.1-TSLPR interface. Structural views compare R90 with the R90S variant, and BLI traces compare TSLPR binding of wild-type TRB5.1 and TRB5.1 R90S. (g) Role of R93 at the predicted TRB5.1-TSLPR interface. Structural views compare R93 with the R93S variant, and BLI traces compare TSLPR binding of wild-type TRB5.1 and TRB5.1 R93S.

From the *de novo* helical bundle strategy, three sorting rounds yielded the top variant TRB2.1, which exhibited a K_D_ of 63.6 nM (R² = 0.998) by BLI, an approximately 29-fold improvement over the parental TRB2.0 **(Fig. 3c)**. From the interface-mimetic strategy, three independent variants, TRB5.1, TRB5.2, and TRB5.3, were isolated from the enriched pool, with K_D_ values of 9.70 nM, 98.2 nM, and 187 nM, respectively **(Fig. 3d, S5a, and S5b)**. The top variant TRB5.1 reached single-digit nanomolar affinity, an approximately 8-fold improvement over the parental TRB5.0. Notably, despite undergoing the same maturation procedure, the matured *de novo* nanobinder (TRB2.1, 63.6 nM) only matched the starting affinity of the interface-mimetic strategy (TRB5.0, 76.8 nM), whereas the matured interface-mimetic nanobinder TRB5.1 attained 9.70 nM, preserving an approximately 7-fold advantage over its *de novo* counterpart. This finding supports the central premise of our two-strategy framework. Grafting a functionally validated natural binding geometry onto a designed scaffold provides a higher attainable affinity ceiling than purely de novo helical scaffolds when comparable maturation resources are invested.

To probe the molecular basis of TSLPR recognition by the matured nanobinders, we focused on TRB5.1 as the highest-affinity variant obtained from our two-strategy design pipeline. We first assessed target selectivity, a critical attribute given that TSLPR belongs to a structurally related family of α-chain cytokine receptors. Binding of TRB5.1 was measured by BLI under identical conditions against a panel of α-chains from the common γ chain (γc) receptor family, namely hIL-2Rα, hIL-4Rα, hIL-7Rα, hIL-9Rα, and hIL-21Rα **(Fig. 3e)**. IL-7Rα was of particular concern because it also serves as the partner chain of TSLPR in the TSLP-TSLPR-IL-7Rα ternary complex. Selective recognition of TSLPR without engaging IL-7Rα was therefore essential to ensure the precision of our heterodimer-disrupting strategy. Quantitative comparison of BLI responses revealed that TRB5.1 produced a clear binding signal only against TSLPR, while responses against all five related interleukin receptors remained at baseline. These results established that TRB5.1 maintains high target selectivity across the structurally related receptor family.

We next examined whether TRB5.1 engages the intended epitope on TSLPR at the molecular level. In the TRB5.1-TSLPR complex model derived from AI-based structural modeling and Rosetta thermodynamic analysis, we identified two arginine residues, R90 and R93, that mediate a network of salt bridges and hydrogen bonds central to the interface and engage complementary residues on the receptor surface. To directly test their functional contributions, we generated single-point variants R90S and R93S and measured their TSLPR binding by BLI **(Fig. 3f and 3g)**. Both substitutions completely abolished the binding signal observed for wild-type TRB5.1. This establishes R90 and R93 as critical recognition residues governing affinity and confirms that TRB5.1 engages TSLPR through specific molecular recognition mediated by the grafted interface-mimetic residues. With these binding and selectivity properties established, we next evaluated all six nanobinders in cell-based assays to determine which variants can functionally antagonize TSLP signaling.

### Cellular Antagonism of TSLP-Mediated Signaling by Interface-Mimetic Nanobinders

To determine whether the six nanobinders from our design and maturation pipeline can block TSLP signaling at the cellular level, we established a reporter assay in HEK293T cells co-transfected with TSLPR and IL-7Rα together with a STAT5-response luciferase reporter (STAT5-Luc). Cells were stimulated with recombinant TSLP and then treated with each nanobinder across a range of concentrations, and STAT5-dependent transcriptional activity was quantified as luciferase activity **(Fig. 4a-4f)**. TSLP stimulation alone induced robust STAT5-Luc activity. The *de novo* binders TRB2.0 and TRB2.1 produced no or only minimal suppression of this signal **(Fig. 4a and 4b)**, and the unmatured interface-mimetic baseline TRB5.0 produced only partial suppression that did not fall below ∼50% **(Fig. 4c)**. In contrast, the three SSM-matured interface-mimetic variants TRB5.1, TRB5.2, and TRB5.3 each produced clear dose-dependent suppression, with TRB5.1 the most potent (p < 0.001; **Fig. 4d-4f**). Notably, TRB5.2 (K_D_ = 98.2 nM) and TRB5.3 (K_D_ = 187 nM) suppressed signaling more completely than the higher-affinity TRB2.1 (K_D_ = 63.6 nM) and TRB5.0 (K_D_ = 76.8 nM), despite binding TSLPR more weakly, showing that target-binding affinity alone does not predict functional antagonism. Because the three most potent variants are all SSM-matured products of the interface-mimetic strategy, an interface-mimetic graft refined by SSM-based maturation appears necessary to convert target binding into complete antagonism of TSLPR-IL-7Rα heterodimer assembly.

**Figure 4.**
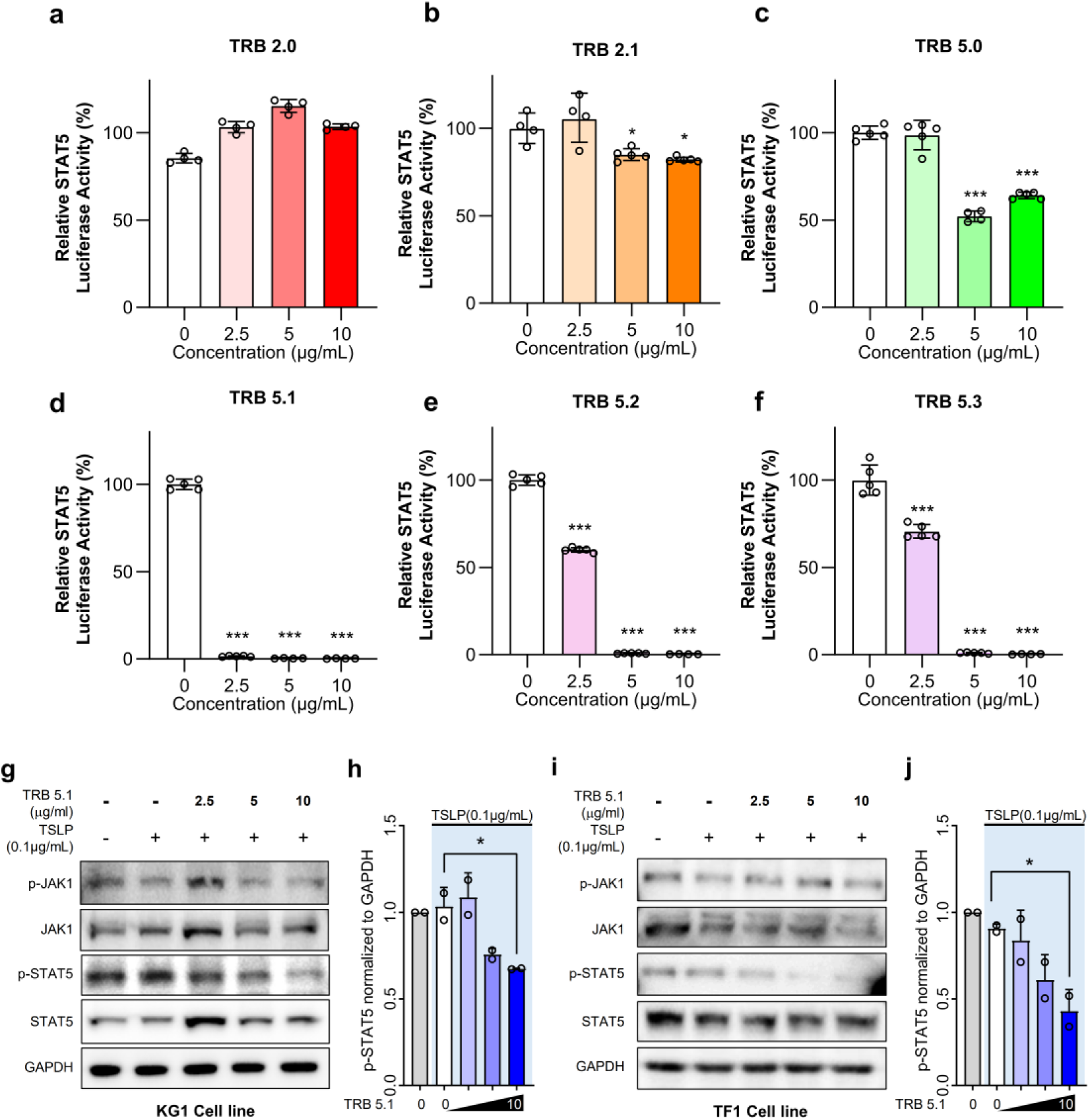
Cellular inhibition assay of TSLP-induced signaling by TSLPR nanobinders. (a-f) Relative STAT5 luciferase activity of HEK293T cells treated with TRB2.0 (a), TRB2.1 (b), TRB5.0 (c), TRB5.1 (d), TRB5.2 (e), or TRB5.3 (f). Luciferase activity (%) is plotted against nanobinder concentration (0, 2.5, 5, 10 μg/mL) for each variant. Data are presented as mean ± SD with individual replicates shown (n = 5). Statistical significance was determined by one-way ANOVA followed by Dunnett’s multiple comparisons test. *p<0.05, ***p < 0.001. (g) Immunoblot analysis of TSLP-induced signaling in KG1 cells. Cells were treated with TSLP and the indicated concentrations of TRB5.1, and phosphorylated and total JAK1 and STAT5 were detected by immunoblotting, with GAPDH as a loading control. (h) Densitometric quantification of p-STAT5 normalized to GAPDH from (g). Data are presented as mean ± SD (n = 3). Statistical significance was determined by one-way ANOVA followed by Dunnett’s multiple comparisons test. *p < 0.05. (i) Immunoblot analysis of TSLP-induced signaling in TF1 cells. Cells were treated with TSLP and the indicated concentrations of TRB5.1, and phosphorylated and total JAK1 and STAT5 were detected by immunoblotting, with GAPDH as a loading control. (j) Densitometric quantification of p-STAT5 normalized to GAPDH from (i). Data are presented as mean ± SD (n = 3). Statistical significance was determined by one-way ANOVA followed by Dunnett’s multiple comparisons test. *p < 0.05.

As TRB5.1 did not affect the viability of KG1 or TF1 cells over the concentration range used in our assays **(Fig. S6)**, we next investigated its effects on endogenous TSLPR signaling. We treated two myeloid leukemia cell lines carrying an endogenous TSLP-TSLPR axis, KG1 and TF1, with TSLP and increasing concentrations of TRB5.1, then analyzed the lysates by immunoblotting. In both lines, TSLP induced strong phosphorylation of JAK1 (p-JAK1) and STAT5 (p-STAT5), and TRB5.1 reduced both signals in a dose-dependent manner **(Fig. 4g and 4i)**. GAPDH levels were unchanged across all lanes, indicating equal loading of total protein. Densitometric analysis of the immunoblots showed that p-STAT5 levels normalized to GAPDH were significantly reduced by higher concentrations of TRB5.1 in both KG1 and TF1 cells **(Fig. 4h and 4j)**. This pattern indicates that TRB5.1 acts specifically at the phosphorylation step rather than by altering target-protein expression. Taken together, although several variants engage TSLPR at nanomolar affinity, only the SSM-matured interface-mimetic variants fully antagonized TSLP-driven signaling, with TRB5.1 most consistently blocking the TSLP-TSLPR axis across three independent cellular models. These results show that combining interface-mimetic motif grafting with SSM-based affinity maturation yields functional nanobinders that disrupt cytokine receptor heterodimer assembly rather than merely binding the target.

### TRB5.1 Suppresses TSLP-Induced Cytokine Gene Expression

Having shown that TRB5.1 suppresses TSLP-induced JAK1 and STAT5 phosphorylation in KG1 and TF1 cells, we next asked whether this blockade extends to the downstream transcriptional output of the pathway. We measured the mRNA levels of three downstream inflammatory targets of TSLP signaling, *IL6*, *CCL17*, and *CCL22*, by RT-qPCR. KG1 and TF1 cells were first stimulated with TSLP, then treated with increasing concentrations of TRB5.1, and transcript levels were normalized to GAPDH. TSLP stimulation alone significantly upregulated all three transcripts in both cell lines, with *CCL22* most strongly induced and *IL6* and *CCL17* induced more modestly. TRB5.1 treatment reversed this induction in a dose-dependent manner, with the highest dose returning *IL6, CCL17*, and *CCL22* toward or below their unstimulated baselines **(Fig. 5a and 5b)**.

**Figure 5.**
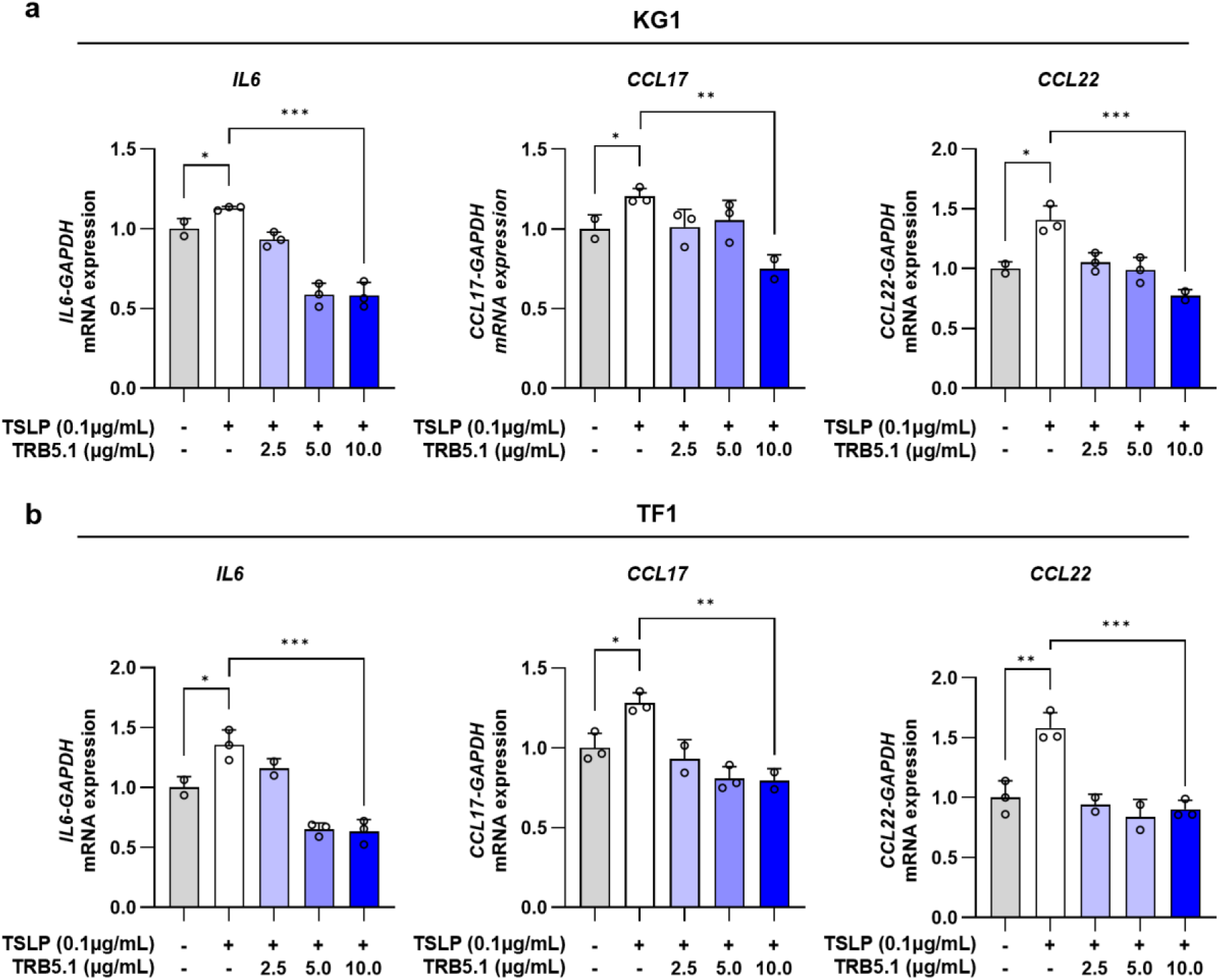
Downstream transcriptional response following TSLP stimulation and nanobinder treatment. (a) RT-qPCR analysis of TSLP-induced inflammatory gene expression in KG1 cells. (b) RT-qPCR analysis of TSLP-induced inflammatory gene expression in TF1 cells. For both (a) and (b), Relative mRNA levels of IL6, CCL17, and CCL22 were quantified and normalized to GAPDH following TSLP stimulation in the presence of increasing concentrations of TRB5.1. Data are presented as mean ± SD with individual replicates shown (n = 3). Statistical significance was determined by one-way ANOVA followed by Dunnett’s multiple comparisons test. *p < 0.05, **p < 0.01, ***p < 0.001.

The same dose-dependent suppression was observed in both KG1 and TF1, two lines carrying an endogenous TSLP-TSLPR axis, confirming that the effect operates on native target genes rather than only on the engineered reporter used in Fig. 4. The transcriptional response also tracked the dose-dependent loss of p-STAT5 measured by immunoblotting in the same cells **(Fig. 4g-4j)**, consistent with transcriptional suppression following directly from the upstream signaling blockade. The three genes differed in sensitivity, with *IL6* falling below its unstimulated level at intermediate doses, whereas *CCL17* and *CCL22* required the highest concentration for significant suppression. Together, these results show that TRB5.1 suppresses not only TSLP-induced JAK-STAT phosphorylation but also the downstream inflammatory transcriptional program in cells with endogenous receptor expression.

### TRB5.1 Broadly Suppresses TSLP-Induced Inflammatory Gene Programs

To determine whether this antagonism extends beyond individual target genes to the broader TSLP-driven program, we profiled the transcriptomes of unstimulated, TSLP-stimulated, and TSLP plus TRB5.1-treated TF1 cells by RNA sequencing. Across 28,395 detected genes, differential expression analysis between the TRB5.1 and TSLP-only conditions identified 135 significantly changed genes, of which 84 were downregulated and 51 upregulated by TRB5.1. The upregulated genes were fewer and of smaller magnitude. The downregulated set contained the major inflammatory effectors of the pathway, with *CXCL1, CXCL8, TNFAIP3, NFKBIA, NFKBIZ,* and *JUN* among the most strongly suppressed transcripts **(Fig. 6a)**.

**Figure 6.**
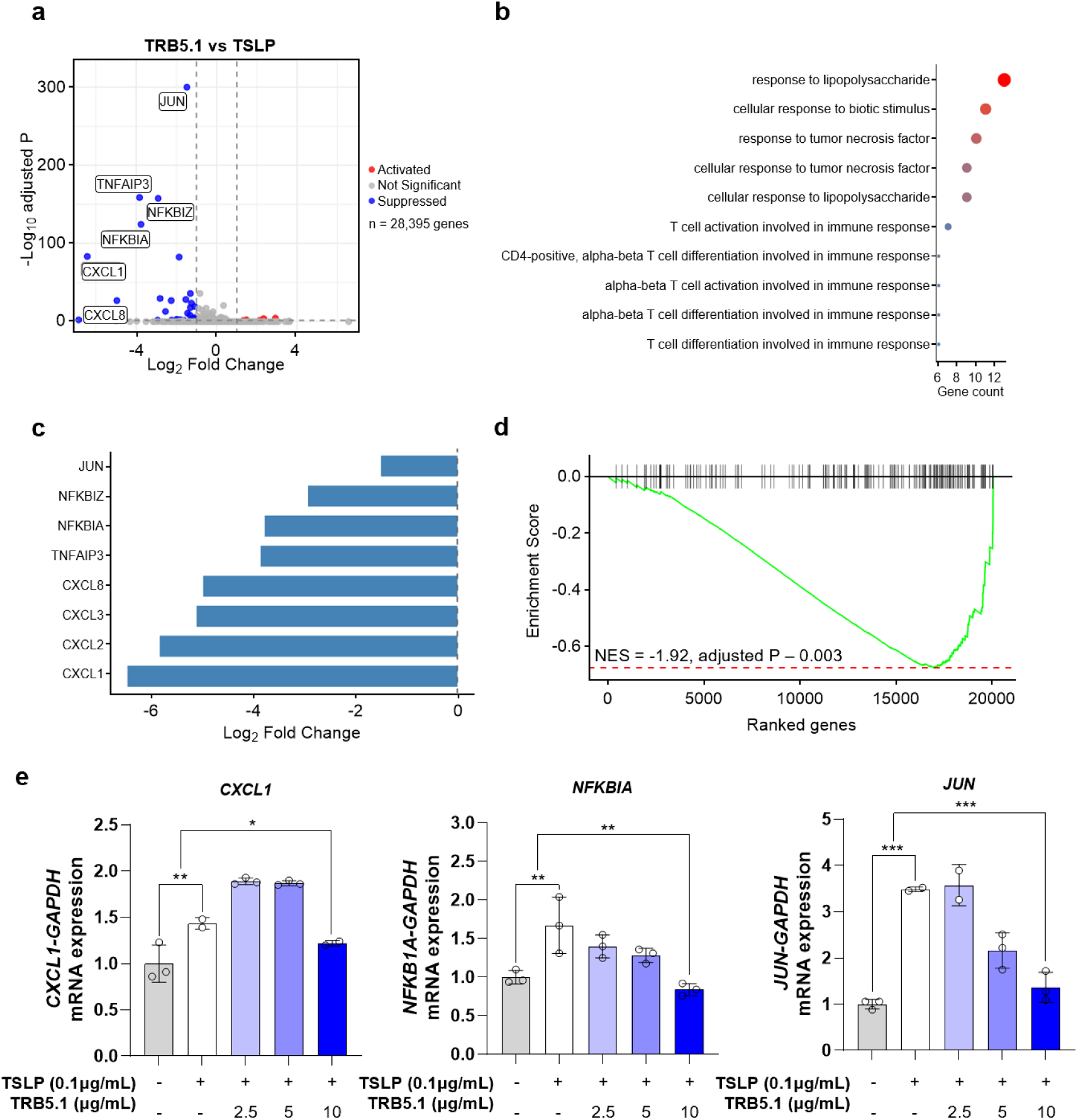
Transcriptomic Suppression of TSLP-Induced Inflammatory Programs by TRB5.1. (a) Volcano plot of differentially expressed genes between TRB5.1-treated (10 μg/mL) and TSLP-stimulated cells. Each point represents one of 28,395 detected genes. The x-axis shows log₂ fold change, and the y-axis shows −log_10_ adjusted P value. Significantly downregulated genes are highlighted in blue and significantly upregulated genes in red, with representative genes (*CXCL1, CXCL8, TNFAIP3, NFKBIA, NFKBIZ, JUN*) labeled. (b) GO Biological Process enrichment analysis of genes downregulated by TRB5.1 treatment. Dot size indicates gene count, and color indicates −log_10_ adjusted P value. (c) Log_2_ fold changes of representative inflammation-related genes suppressed by TRB5.1 relative to TSLP stimulation. CXCL-family chemokines (*CXCL1, CXCL2, CXCL3, CXCL8*) and NF-κB-associated genes (*TNFAIP3, NFKBIA, NFKBIZ, JUN*) are shown. (d) GSEA enrichment plot for the Hallmark TNF-α Signaling via NF-κB gene set (NES = −1.92, adjusted P = 0.003). Running enrichment score curves are shown above the ranked gene list, with black ticks indicating positions of gene set members. (e) RT-qPCR validation of representative RNA-seq-identified inflammatory genes (*CXCL1, NFKBIA, JUN*) in TF-1 cells pretreated with TRB5.1 (2.5, 5, 10 μg/mL) before TSLP stimulation (0.1 μg/mL). mRNA levels were normalized to *GAPDH*. Data are mean ± SD (n = 3, individual replicates shown). One-way ANOVA with Dunnett’s test. *P < 0.05, **P < 0.01, ***P < 0.001.

Gene Ontology (GO) analysis of the downregulated genes resolved into two functional groups. The first comprised innate inflammatory processes, including response to lipopolysaccharide, response to tumor necrosis factor, cellular response to tumor necrosis factor, and cellular response to biotic stimulus. The second comprised adaptive programs, including T cell activation and CD4-positive alpha-beta T cell differentiation **(Fig. 6b)**. Response to lipopolysaccharide was the most significantly enriched term and carried the highest gene count. Inspecting the genes behind these terms, the CXCL-family chemokines *CXCL1, CXCL2, CXCL3,* and *CXCL8* showed the largest reductions, exceeding five log₂ units for *CXCL1* and *CXCL2*. The NF-κB-associated genes *TNFAIP3, NFKBIA, NFKBIZ*, and *JUN* were suppressed to a smaller but consistent degree **(Fig. 6c)**.

To test whether these changes reflected coordinated movement of a defined inflammatory program rather than isolated genes, we performed gene set enrichment analysis against the Hallmark collection. TNF-α signaling via NF-κB showed significant negative enrichment (NES = −1.92, adjusted P = 0.003), with the running score reaching a minimum near −0.65 and the leading edge concentrated among the most strongly downregulated genes **(Fig. 6d)**. The inflammatory response gene set was likewise negatively enriched (NES = −1.56, adjusted P = 0.010). Principal componen analysis separated the three conditions, with the first component aligning with the TSLP response. Along this axis the TSLP plus TRB5.1 samples sat between the TSLP-only and unstimulated samples, indicating a partial return toward the resting state **(Fig. S7)**.

To confirm these transcriptome-level changes by an independent method, we measured three representative genes from the suppressed TNF-α/NF-κB inflammatory program, namely the chemokine *CXCL1*, the NF-κB feedback inhibitor *NFKBIA*, and the AP-1 component *JUN*, by RT-qPCR in TF1 cells, the same line profiled by RNA sequencing. Cells were pretreated with increasing concentrations of TRB5.1 and stimulated with TSLP, and transcript levels were normalized to *GAPDH*. TSLP alone significantly upregulated all three transcripts relative to the unstimulated control, with *JUN* induced most strongly and *CXCL1* and *NFKBIA* induced more modestly. At the highest concentration of TRB5.1, all three transcripts were significantly reduced relative to the TSLP-only condition, with *NFKBIA* and *JUN* declining progressively across the concentration range and *CXCL1* reaching significant suppression only at the highest concentration **(Fig. 6e)**. Together, these results show that TRB5.1 suppresses not only individual target genes but the broader NF-κB-associated inflammatory transcriptional network activated by TSLP, a conclusion supported by both transcriptome-wide RNA sequencing and independent RT-qPCR validation of representative genes.

## Discussion

This study establishes TRB5.1 as, to our knowledge, the first *de novo* nanobinder to antagonize TSLP-TSLPR signaling by sterically occluding assembly of the TSLPR-IL-7Rα heterodimer. By evaluating two structure-guided strategies against a single target in parallel, the work also provides a controlled comparison of design principles for expansive, water-mediated cytokine-receptor interfaces. Such interfaces depend on distributed networks of weak polar contacts that a purely *de novo* scaffold reproduces only at the expense of stability. Reusing the binding geometry validated in the native complex therefore proved more effective than unconstrained helical design. We grafted the two TSLP contact elements, the α4 helix and the binding loop, onto a hyper-stable *de novo* scaffold and removed the disordered loop. This preserved the native recognition surface and raised the melting temperature to 97.35 °C (TRB5.0), roughly 40 °C above the *de novo* binder (TRB2.0, 57.15 °C). After maturation, the lead TRB5.1 reached single-digit nanomolar affinity (K_D_ = 9.70 nM), whereas the matured *de novo* binder reached only 63.6 nM (TRB2.1). Purely helical design did not attain this combination of affinity and thermal stability.

A central mechanistic finding is the decoupling of target-binding affinity from functional antagonism. Following site-saturation maturation, several interface-mimetic variants of modest affinity, notably TRB5.2 (K_D_ = 98.2 nM) and TRB5.3 (K_D_ = 187 nM), suppressed STAT5 signaling more completely than the tighter-binding TRB2.1 (K_D_ = 63.6 nM) and the unmatured TRB5.0 (K_D_ = 76.8 nM). This indicates that affinity alone does not predict cellular potency. The ternary assembly mechanism accounts for the decoupling. TSLP first engages TSLPR and induces a conformational change that exposes the surface required for IL-7Rα recruitment [6]. Functional antagonism therefore demands occupancy of the receptor in a geometry that sterically precludes this transition, not mere engagement of the cleft. We propose that the rigid surfaces of the *de novo* binders, despite high-affinity occupancy, leave TSLPR conformationally permissive for IL-7Rα docking. The grafted native loop of TRB5.1, in contrast, acts as a structural wedge that constrains the receptor in an assembly-incompetent conformation. For multi-subunit receptor systems, design should therefore target geometric and steric preclusion rather than the maximization of binding energy alone.

Therapeutically, the position of TRB5.1 within the pathway confers a distinct advantage, as corroborated by transcriptome-wide profiling. TSLP acts as an upstream regulator at the apex of the type 2 inflammatory cascade. Antagonism at its receptor therefore produced a coordinated reversal of the downstream program rather than suppression of a single effector arm. Across 28,395 detected genes, RNA sequencing identified 135 differentially expressed genes and revealed broad downregulation of the NF-κB and TNF-α-associated network, from chemokines such as *CXCL1* and *CXCL8* to the NF-κB regulators *NFKBIA* and *NFKBIZ*. RT-qPCR independently confirmed suppression of downstream type 2 chemokines including *CCL17* and *CCL22*. Gene set enrichment confirmed significant depletion of TNF-α signaling via NF-κB. This program-level suppression is an advantage over agents that neutralize individual downstream effectors, such as dupilumab against IL-4Rα or mepolizumab against IL-5 [26–28]. The strict selectivity of TRB5.1, which showed no binding to five structurally related γc-family α-chain receptors, confines its activity to the TSLP axis and avoids the broad immunosuppression associated with pan-JAK inhibitors or systemic corticosteroids.

TRB5.1 also differs mechanistically from tezepelumab, the approved 150 kDa monoclonal antibody that sequesters soluble TSLP and must be maintained in excess during cytokine surges [8]. By targeting the finite and steady-state surface receptor pool instead, TRB5.1 can approach saturation largely independently of cytokine output. Its compact size (∼10 kDa) and thermal resilience further render it amenable to aerosolized or topical administration. Clinical translation will nonetheless require *in vivo* efficacy, pharmacokinetic, and immunogenicity studies, since a protein well below the ∼60 to 70 kDa glomerular filtration threshold is subject to rapid renal clearance and to anti-drug antibody formation. Systemic indications would additionally require half-life extension. More broadly, this work reframes disruption of a pathogenic cytokine-receptor heterodimer as a geometrically constrained design objective. It thereby provides a scalable blueprint and positions TRB5.1 as both a developable therapeutic lead and a template for the broader shared-chain cytokine network.

## Materials and methods

### *In Silico* Design of TSLPR-Binding Nanobinders

The human TSLP/TSLPR/IL-7Rα ternary complex (PDB ID 5J11) was used as the structural template. Spatial Aggregation Propensity (SAP) scoring of the TSLPR surface in PyMOL (Schrödinger) localized a hydrophobic patch centered on I66, W84, and Y87, which was defined as the design hot-spot. Two complementary structure-guided strategies were then implemented within a unified RFdiffusion–ProteinMPNN–AlphaFold3 (AF3) pipeline [16–18]. In the *de novo* helical bundle strategy, backbones of 70–80 residues targeting the I66/W84/Y87 hot-spot were generated with RFdiffusion. From 100 initial designs, 16 compact helical bundles were selected; each was diversified by partial diffusion (20 variants per backbone), and two sequences per variant were generated with ProteinMPNN. Complexes with the TSLPR extracellular domain were predicted by AF3, and models with ipTM ≥ 0.85 were iteratively refined by Rosetta Relax or further partial diffusion, redesigned with ProteinMPNN, and re-predicted until interface confidence converged. 24 final candidates were prioritized by combined pLDDT and interchain pAE filtering. In the interface-mimetic grafting strategy, two TSLPR-binding elements of human TSLP, the loop (residues 53-74) and the α4 helix (residues 139-159), were extracted from PDB 5J11 and fixed as binding motifs. RFdiffusion was used to design four-helix-bundle scaffolds that rigidly display both motifs in their native orientation, and non-motif positions were designed with ProteinMPNN. AF3-predicted complexes with ipTM ≥ 0.85 were refined by Rosetta Relax, redesigned with ProteinMPNN, and re-predicted iteratively. From the resulting pool of 100 sequences, 28 candidates were prioritized by interchain pAE and pLDDT.

### Yeast Surface Display Screening

Designed binder genes (Integrated DNA Technologies) were co-transformed with linearized pETCON3 into *Saccharomyces cerevisiae* EBY100 by electroporation. Transformants were grown in SDCAA (30 °C, 48 h), induced in SGCAA at OD₆₀₀ = 1.0 (20 °C, 36 h), washed in PBSA (PBS + 1% BSA), and co-stained with PE-conjugated anti-Myc antibody (1:50; Cell Signaling Technology) and 500 nM His-tagged TSLPR pre-complexed with anti-His-FITC antibody (Cell Signaling Technology) for 30 min at 4 °C. Double-positive cells were isolated on a CytoFLEX SRT cell sorter (Beckman Coulter) and recovered in SDCAA. Plasmids were extracted (Zymoprep Yeast Plasmid Miniprep II; Zymo Research), transformed into *E. coli* DH10β, and individual clones were confirmed by Sanger sequencing.

### Affinity Maturation by Site Saturation Mutagenesis (SSM)

Interface residues of TRB2.0 and TRB5.0 were identified from the AF3 binder–TSLPR complexes using the PDBePISA server. Each interface position was diversified using NNK-degenerate primers (Integrated DNA Technologies), and SSM libraries were assembled by overlap-extension PCR. Libraries were displayed and sorted as above under increasingly stringent TSLPR concentrations (500, 100, 20 nM) over successive rounds, and enriched variants were identified by Sanger sequencing.

### Protein Expression and Purification

Nanobinders were expressed as N-terminal His₆–MBP–TEV fusions in *E. coli* BL21(DE3). Cultures in LB were induced at OD₆₀₀ = 0.6-0.8 with 0.5 mM IPTG and grown at 18 °C for 18 h. Cells were lysed by sonication in 20 mM Tris-HCl pH 7.5, 500 mM NaCl, 25 mM imidazole, and clarified lysates were applied to a HisTrap HP 5 mL column (Cytiva). Eluted fusion protein was cleaved with TEV protease (1:30 molar ratio) during overnight dialysis (4 °C), reapplied to HisTrap HP, and the untagged binder was collected in the flow-through. Final polishing was performed on a HiLoad 26/600 Superdex 75 pg column (Cytiva) in DPBS. The final purity of the nanobinders was assessed by SDS-PAGE **(Fig. S8)**. Monomeric fractions were concentrated to 9.7 mg/mL(10 kDa MWCO; Merck Millipore) and stored at −80 °C.

### SEC-MALS and DSC

SEC-MALS was performed at the Korea Basic Science Institute (KBSI) using a HELEOS detector with an rEX refractometer (Wyatt Technology) at 0.5 mL/min, and data were analyzed in ASTRA 6 (Zimm fit, dn/dc = 0.1850 mL/g). Thermal stability was measured on a MicroCal PEAQ-DSC (Malvern) at 122 µM in DPBS, scanned 20-120 °C at 1 °C/min; Tm was determined from the Cp peak using MicroCal PEAQ-DSC software v1.64.

### Bio-Layer Interferometry (BLI)

Binding kinetics were measured on an Octet R8 system (Sartorius) with SA biosensors. Biotinylated TSLPR-Fc-Avi (3 ng/µL; AcroBiosystems) was loaded for 300 s, followed by 300 s association with nanobinders at 7.8-500 nM and 600 s dissociation in kinetics buffer (PBS pH 7.4, 0.01% BSA, 0.002% Tween-20). Sensorgrams were referenced and globally fitted to a 1:1 binding model in Octet Analysis Studio v12.2.2.26 (Sartorius); fits with R² > 0.95 were accepted. Specificity was tested against human IL-2Rα, IL-4Rα, IL-7Rα, IL-9Rα, and IL-21Rα under identical conditions, and TRB5.1(R90S) and TRB5.1(R93S) served as interface knockout controls.

### Cell Culture

KG1 and TF1 cells were maintained in RPMI-1640 (Thermo Fisher Scientific) with 20% FBS (Gibco) and 1% penicillin-streptomycin; HEK293T cells were cultured in DMEM with 10% FBS and 1% penicillin-streptomycin. All cells were maintained at 37 °C, 5% CO₂.

### STAT5-Luciferase Reporter Assay

HEK293T cells (5 × 10⁵ per well, 6-well plate) were co-transfected with pMET7-Flag-TSLPR (400 ng), pMET7-HA-IL7Rα (400 ng), and pGL4.52-STAT5 (200 ng) using FuGENE® HD (Promega). After 24 h, nanobinders (2.5, 5, 10 µg/mL) were added for 2 h, followed by 0.1 µg/mL recombinant human TSLP (PeproTech) for 24 h. Luciferase activity was measured using the Luciferase Assay System (Promega) according to the manufacturer’s instructions.

### Western Blot

KG1 and TF1 cells were pretreated with TRB5.1 (2.5, 5, 10 µg/mL, 2 h) and stimulated with TSLP (0.1 µg/mL, 1 h). Lysates in RIPA buffer (LPS Solution) supplemented with protease/phosphatase inhibitors (Thermo Fisher Scientific) were quantified by BCA assay (Thermo Fisher Scientific). Proteins (30 µg) were resolved on 8% SDS-PAGE, transferred to PVDF (MilliporeSigma), blocked in 3% BSA, and probed with antibodies against p-JAK1 (#3331), p-STAT5 (#9351), JAK1 (#3344), STAT5 (#25656) (Cell Signaling Technology), and GAPDH (sc-47724; Santa Cruz). HRP-conjugated secondaries (#7074S, #7076S; Cell Signaling Technology) were detected by ECL (Cytiva).

### RT-qPCR

Total RNA was extracted using TRIzol (Thermo Fisher Scientific), reverse-transcribed with iScript™ (Bio-Rad), and amplified with iQ™ SYBR® Green Supermix (Bio-Rad) using primers for *IL6, CCL17, CCL22, CXCL1, NFKBIA, JUN,* and *GAPDH*. Relative expression was calculated by the comparative Ct (2^−ΔΔCt) method with *GAPDH* normalization.

### RNA Sequencing & Statistical Analysis

Total RNA from unstimulated, TSLP-stimulated, and TRB5.1+TSLP-treated cells was processed with the TruSeq Stranded mRNA Library Prep Kit (Illumina) and sequenced on an Illumina NovaSeq 6000 (paired-end 150 bp). Reads were aligned to GRCh38 with STAR, quantified with featureCounts, and analyzed for differential expression with DESeq2 (|log₂FC| ≥ 1, adjusted P < 0.05) Differential expression and downstream analyses focused primarily on genes differentially expressed between TSLP-stimulated and TRB5.1-pretreated TSLP-stimulated samples. PCA, Gene Ontology (GO) Biological Process enrichment, and Gene Set Enrichment Analysis (GSEA) using MSigDB Hallmark gene sets were performed in R (version 4.5.3)

## CRediT authorship contribution statement

Woo-Chan Ahn: Writing – review & editing, Writing – original draft, Software, Methodology, Investigation. Min-Jung Son: Investigation, Formal analysis, Data curation. Jae-Rin Kim: Validation, Methodology, Investigation. Seong-Ryeong Go: Methodology, Investigation. Ju-Eun Kang: Investigation. Junsoo Park: Investigation. Young-Hoon Lee: Investigation. Su-Jin Lee: Investigation. Jeong-Hoon Kim: Supervision, Methodology. Juyeon Jung: Supervision, Methodology. Eui-Jeon Woo: Writing – review & editing, Supervision, Project administration, Funding acquisition, Conceptualization. Young-Joo Jeon: Writing – review & editing, Supervision, Investigation, Funding acquisition, Formal analysis, Conceptualization. Kwang-Hyun Park: Writing – review & editing, Writing – original draft, Supervision, Software, Project administration, Methodology, Funding acquisition, Conceptualization.

## Declaration of Generative AI and AI-assisted technologies in the writing process

During the preparation of this work, the authors used ChatGPT 5 to edit grammatical errors. After using this tool/service, the authors reviewed and edited the content as needed and take full responsibility for the content of the published article.

## Data availability

The datasets supporting the conclusions of this article are included within the article and its additional files. All original data and information on reagents and protocols are available upon request. Correspondence and requests for materials should be addressed to Kwang-Hyun Park.

## Acknowledgments

This work was supported by National Research Foundation of Korea (NRF) grants funded by the Korean Government (MSIP) (RS-2022-NR071772, RS-2021-NR059435 and 2021M3A9G802559922). This research was partially supported by the National Research Council of Science & Technology (NST) (CRC22025-500), and the KRIBB Research Initiative Program (KGM1062612, KGM5382632, KQM0052611 and KGM1322612).

